# Complexity of Brain Activity and Connectivity in Functional Neuroimaging

**DOI:** 10.1101/377028

**Authors:** Stavros I. Dimitriadis

## Abstract

Understanding the complexity of human brain dynamics and brain connectivity across the repertoire of functional neuroimaging and various conditions, is of paramount importance. Novel measures should be designed tailored to the input focusing on multichannel activity and dynamic functional brain connectivity (DFBC).

Here, we defined a novel complexity index (CI) from the field of symbolic dynamics that quantifies patterns of different words up to a length from a symbolic sequence. The CI characterizes the complexity of the brain activity.

We analysed dynamic functional brain connectivity by adopting the sliding window approach using imaginary part of phase locking value (iPLV) for EEG/ECoG/MEG and wavelet coherence (WC) for fMRI. Both intra and cross-frequency couplings (CFC) namely phase-to-amplitude were estimated using iPLV/WC at every snapshot of the DFBC. Using proper surrogate analysis, we defined the dominant intrinsic coupling mode (DICM) per pair of regions-of-interest (ROI). The spatio-temporal probability distribution of DICM were reported to reveal the most prominent coupling modes per condition and modality. Finally, a novel flexibility index is defined that quantifies the transition of DICM per pair of ROIs between consecutive time-windows.

The whole methodology was demonstrated using four neuroimaging datasets (EEG/ECoG/MEG/fMRI).

Finally, we succeeded to totally discriminate healthy controls from schizophrenic using FI and dynamic reconfiguration of DICM. Anesthesia independently of the drug caused a global decreased of complexity in all frequency bands with the exception in δ and alters the dynamic reconfiguration of DICM. CI and DICM of MEG/fMRI resting-state recordings in two spatial scales were high reliable.

**Significant Statement:** In the present study, we demonstrated novel indexes for the estimation of complexity in both raw brain activity and dynamic functional connectivity. To ort the universality of both indexes for the majority of functional neuroimaging modalities, we adopted open datasets from electro and magneto-encephalography, from electro-corticography and functional magnetic resonance imaging. Both indexes proved informative and reliable across repeat scan sessions. Moreover, we succeeded to totally discriminate with absolute accuracy healthy controls from schizophrenic patients. Both indexes proved sensitive to common anesthetic drugs effect in monkeys and reliable in MEG and fMRI repeat scan sessions. We first reported the notion of cross-frequency coupling in BOLD activity. Our analysis could be adapted it in any task and modality for any hypothesis driven study.

## 1. Introduction

There are many complex systems in nature that change over time where their functionality could be estimated by applying traditional methods to the recorded time series. Especially, in the case that the time series that characterize the complex system are simple and linear then simple approaches like Fourier transform can characterize the signal patterns. More complex systems such as chaotic oscillations, bifurcations and more challenging brain activity demand more sophisticated approaches that deal with the metastability and non-linearity of the underlying functionality (Gao et al., 2011; Dimitriadis et al., 2012).

One of the most important family of techniques in temporal data mining is the symbolic time-series analysis. The general idea of symbolization is the transformation of a raw time-series into a sequence of discrete symbols. The whole approach opens a variety of available neuroinformatic tools that share common theoretical background with Markov chain (Seneta, 1981), bioinformatics (Baldi and Brunak, 1998) and with theory of communication (Shannon and Weaver, 1998).

A well-know method for analyzing the time-delay embedding of a 1D time series time-series that describes a dynamical system is the recurrence plots (Marwan et al., 2007). This method reveals the dynamical invariants of a system. A new technique called Fuzzy Symbolic Dynamics (FSD) clusters each multidimensional point to a neighborhood and then maps this information to a simplified two or three dimensional diagram (Duch and Dobosz 2011).

A large set of complexity estimators derived from information theory have been applied to symbolic sequences from multichannel EEG/MEG data in order to further understand brain dynamics, to design novel diagnostic tools tailored to brain diseases and also to discriminate brain activity between different cognitive tasks (Gao et al., 2011). A notable attention has been given to Lempel-Ziv Complexity (LZ) which quantifies different substrings in the binarized symbolic time series (Lempel-Ziv, 1976). The binarization threshold is the mean amplitude of the time-series in most of the cases. LZ complexity has been used in recordings from rats during sleep (Abasolo et al., 2015), during propofol anesthesia (Hudetz et al., 2016), in Alzheimer’s disease (Abasolo et al., 2006), in traumatic brain injury (Luo et al., 2013), in dyslexia (Dimitriadis et al., 2017), in mild traumatic brain injury (Antonakakis et al., 2017) etc.

LZ complexity has many limitations. First of all, you have to apply an arbitrary threshold scheme that transform the original time series into a symbolic sequence of 0s and 1s. Secondly, the binarization is applied to the 1D time series and not a reconstructed phase space via time-delay embedding. This well-known approach can reveal the nonlinear and complex nature of the system described via the time series.

In two recent studies, we introduced a novel complexity index (CI) based on the transformation of a reconstructed phase space of a 1D time-series via time-delay embedding procedure to a symbolic sequence via neural-gas algorithm (Antonakakis et al., 2017; Dimitriadis et al., 2017). Neural-gas algorithm learns the manifold of the trajectory and transforms it into a symbolic sequence. Then, CI is defined as the distribution of distinct words up to a specific length of letters-symbols normalized by the distribution of distinct words of a number of randomized versions of symbolic time series. We reported higher CI values for dyslexic children versus non-impaired readers using MEG resting-state (Dimitriadis et al., 2017) and lower CI values for mTBI subjects versus healthy controls using MEG resting-state (Antonakakis et al., 2017). CI outperforms LZ complexity after applying machine learning approach for differentiate non-impaired readers from dyslexic children and healthy controls with age-matched mTBI subjects.

Brain dynamics recorded via EEG/MEG are complex signals that encapsulate the activity in various frequencies called brain rhythms. Additionally to the power spectrum analysis of signal as 1D time-series using Fourier or wavelet transform or to more sophisticated analysis such as our CI, it is more than important to reveal the different type of functional interactions. There are two complementary functional coupling mechanisms in spontaneous activity and also in task-related activity, the phase coupling and the coupling of signal envelopes in predefined band-pass filtered brain signals. Both types of intrinsic coupling modes (ICMs) have demonstrated different degree of sensitivity in normal and disease brain activity and also different correlation levels with structural connectivity (Engel et al., 2013). Apart from studying ICMs by taking into account the functional coupling in both amplitude and phase domain between signals with the same frequency content, it is significant to explore also their cross-frequency coupling mechanisms (Jirsa and Müller, 2013; Dimitriadis et al., 2015ba,b,2016a,c,2017).

It is well known that human brain mechanisms support functionally seperated temporal frames to group brain activity into sequences of neural assemblies where multi-frequency interactions simultaneously synchronized across the whole brain. These multiplex interactions create the syntactic rules which are significant for the exchange of information across the cortex (Buzsaki and Watscon, 2012).

Several studies have explored and revealed the physiological significant role of crossfrequency coupling mechanisms. To give an example, the strength of θ-γ coupling in the hippocampus and striatum of the rat was influenced by task demands (Tort et al., 2008, 2009). In another experiment, the coupling strength between a θ (4-Hz) oscillation and *y* power within the prefrontal cortex increased during the working memory phase of a choice task (Fujisawa and Buzsaki, 2011).

The observed phenomenon of cross-frequency coupling mechanism supports the hierarchical organization of multiple brain oscillations across space and time and untangled the simultaneously brain interactions across multiple time scales (Canolty and Knight, 2010; Fell and Axmacher, 2011). Well established computational models have explored the theoretical advantages of the existence of cross-frequency coupling (Lisman and Idiart, 1995; Neymotin et al., 2011). These models revealed the key mechanisms of cross-frequency coupling which may serve as the backbone of a neural syntax. The exist syntactic rules allow for both segmentation of spike trains into cell assemblies (“letters”) and assembly sequences (neural “words”) (Buszaki, 2010).

From the aforementioned evidences, it is more than important to explore the repertoire of available intra and cross-frequency interactions among brain rhythms and between brain areas under the same graph model.

Here, we provided a framework of how to study the majority of available and well established interactions simultaneously and across the whole brain. It is important to explore brain interactions globally and afterward to focus on sub-networks and local interactions. On the top of it, we define an index that quantifies the flexibility of a pair of ROIs which quantifies the exchange rate of the preferred (dominant) coupling mode.

The complexity and the multiplexity of the human brain functional connectivity can be explored on the original functional dimensions only if all the possible interactions are studying under the same framework. For that reason, it is important to study the dominant type of interactions between every pair of brain areas at every snapshot of dynamic functional connectivity graph (DFCG) across the experimental time. We hypothesize that the dynamic reconfiguration of dominant intrinsic coupling modes (DICM) can capture the complexity – multiplexity of human and also the non-human brain during spontaneous activity and also in cognitive tasks. Additionally, the flexibility of this reconfiguration can be directly be linked to the multiplexity of brain functionality and its ability to adjust fast to any environmental stimulus (Buzsáki and Draguhn, 2004; Buzsaki, 2010; Buzsaki et al., 2013). We recently demonstrated the effectiveness of FI via the definition of DICM using a life-span EEG dataset in order to design a chronnectomic Brain Aged Index (CBAI) (Dimitriadis et al., 2017).

Here, we demonstrated the effectiveness of both complexity approaches in raw activity and dynamic functional connectivity using the following four datasets:

1. A EEG study with healthy control and schizophrenic patients at resting-state
2. A ECoG study from a monkey recorded during alter and after anesthesia
3. A MEG repeat-scan study at resting-state
4. A fMRI repeat-scan study from a single subject

In Materials and Methods section, we described the data acquisition and details of the four datasets, the preprocessing steps including the denoising with ICA and the beamforming analysis to reconstruct the sources. The Results section is devoted to describe the results including classification results, significant changes between conditions and reliability of the proposed indexes in repeat scan sessions. Finally, the Discussion section includes the discussion of the current research results with future extensions.

## 2. Materials and Methods

In this section, we describe the datasets including subjects and data acquisition from the four functional neuroimaging modalities. We also described the preprocessing steps and how complexity indexes have been estimated.

### 2.1.1 EEG recordings

The subjects were adolescents who had been screened by psychiatrist and devided into two groups: healthy (n = 39) and with symptoms of schizophrenia (n = 45). Both groups included only school boys. The age of the patients ranged from 10 years and 8 months to 14 years. The control group included 39 healthy schoolboys aged from 11 years to 13 years and 9 months. The mean age in both groups was 12 years and 3 months.

EEG activity was recorded from 16 EEG channels where their electrode positions is demonstrated in Fig.1. The sampling rate is 128 Hz and the recording time was 1 min, thus a total of 7680 samples refer to 1 minute of EEG record. You can download the EEG recordings from the website: http://brain.bio.msu.ru/eeg_schizophrenia.htm. The dataset has been adopted from previous published papers.

**Figure 1.**
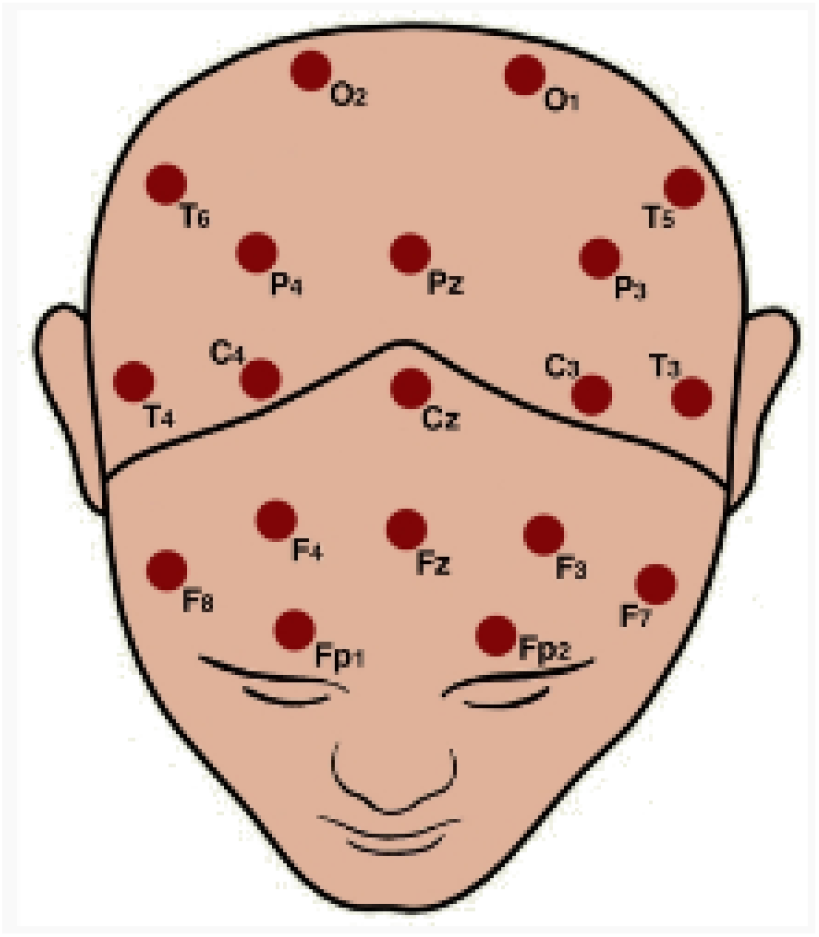
The topographical positions of EEG channels.

### 2.1.2 ECoG Recordings

Chronically implanted, customized multichannel ECoG electrode arrays (Unique Medical, Japan) were used for neural recording (Nagasaka et al., 2011). The array was implanted in the subdural space in 4 adult macaque monkeys (M1-M3 are *Macaca fuscata* and M4 is *Macaca mulatto).* One hundred and twenty-eight ECoG electrodes with an inter-electrode distance of 5 mm were implanted in the left hemisphere, continuously covering over the frontal, parietal, temporal, and occipital lobes. Fig.2 illustrates the positions of the ECoG electrodes. Parts of the dataset are shared in the public server Neurotycho.org (http://neurotycho.org/) . I analysed EcoG recordings from one monkey was sitting calm with head and arm restrained. ECoG data were recorded first with alert and later with anesthetic condition after injecting propofol (Yanagawa et al., 2013). For further details see the original article.

**Figure 2.**
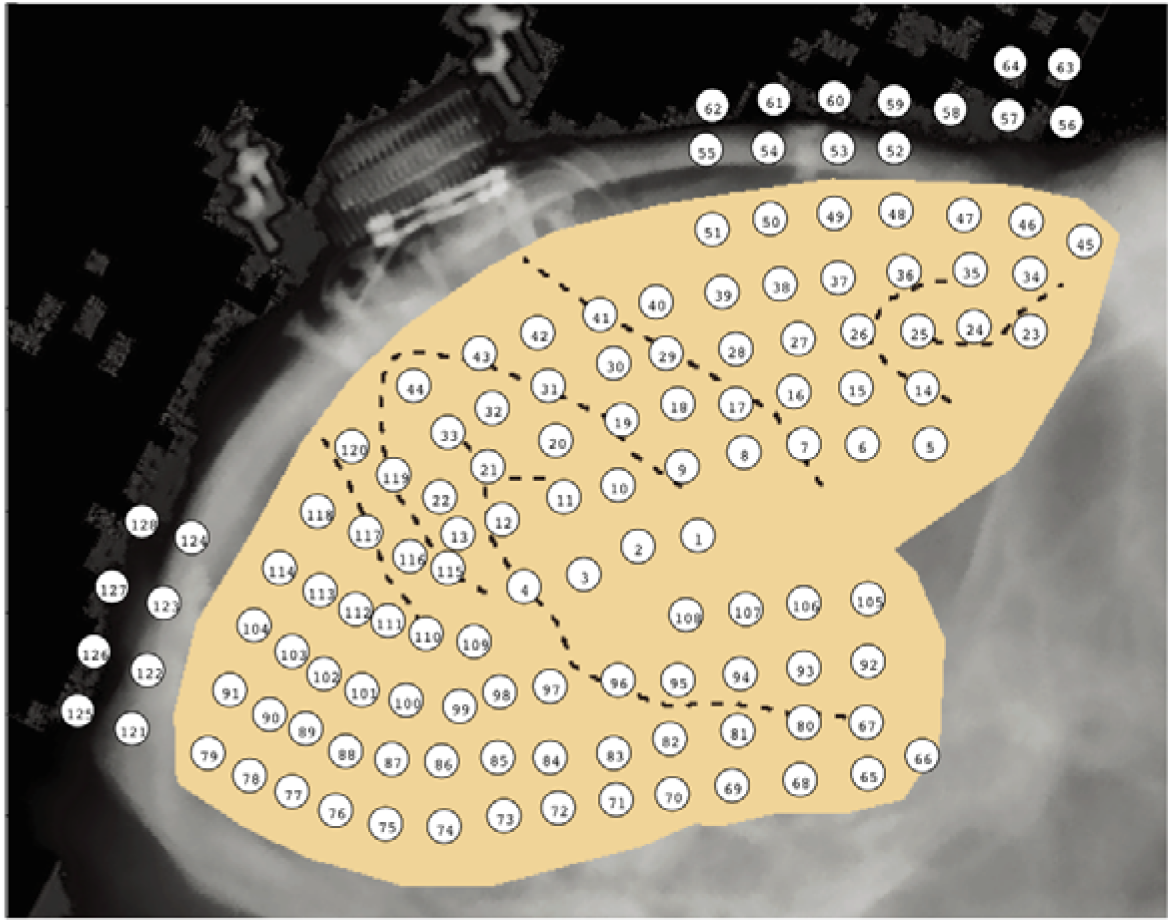
The topographical positions of EcoG channels^1^.

ECoG electrodes have been clustered to the following brain areas:

Medial prefrontal cortex, lateral prefrontal cortex, premotor cortex, primary and somatosensory cortices, parietal cortex, temporal cortex, higher visual cortex and lower visual cortex (see Fig.1 in Yanagawa et al., 2013).

### 2.1.3 MEG Recordings

#### 2.1.3.1 Subjects

40 healthy subjects (age *22.85*}*3.74*years, 15 women and 25 men) underwent two resting-state MEG sessions (eyes open) with a 1-week test-retest interval. For each participant, scans were scheduled at the same day of the week and same time of the day. The duration of MEG resting-state was 5 mins for every participant. The study was approved by the Ethics Committee of the School of Psychology at Cardiff University, and participants provided informed and written consent.

#### 2.1.3.2 MEG-MRI Recordings

Whole-head MEG recordings were made using a 275-channel CTF radial gradiometer system. 29 reference channels were recorded for noise cancellation purposes and the primary sensors were analysed as synthetic third-order gradiometers (Vrba and Robinson, 2001). Two or three of the 275 channels were turned off due to excessive sensor noise (depending on time of acquisition). Subjects were seated upright in the magnetically shielded room. To achieve MRI/MEG co-registration, fiduciary markers were placed at fixed distances from three anatomical landmarks identifiable in the subject’s anatomical MRI, and their locations were verified afterwards using high-resolution digital photographs. Head localisation was performed before and after each recording, and a trigger was sent to the acquisition computer at relevant stimulus events. For further details see Dimitriadis et al., 2018.

### 2.1.4 fMRI Single-Case Long Term Dataset

The participant on this single-case study (author R.A.P.) is a right-handed Caucasian male, aged 45 years at the onset of the study. RS-fMRI was recorded in one hundred scans throughout the data collection period (89 in the production phase), using a multi-band EPI sequence (TR = 1.16 ms, TE = 30 ms, flip angle = 63 degrees (the Ernst angle for gray matter), voxel size = 2.4 × 2.4 × 2 mm, distance factor = 20%, 68 slices, oriented 30 degrees back from AC/PC, 96 × 96 matrix, 230 mm FOV, MB factor = 4, 10:00 scan length).

### 2.2 Preprocessing Steps

In this section, we described the denoising step of brain activity for EEG/MEG/ECoG/fMRI datasets, the beamformer analysis for MEG dataset and the filtering set up on predefined frequency bands.

### 2.2.1 Independent Component Analysis

Ongoing activity from each modality was corrected for artifacts through the following procedure. Line noise was removed with a notch filter at 60 Hz and the data recording from a single subject was whitened and reduced in dimensionality by means of Principal Component Analysis (PCA) with a threshold corresponding to 95% of total variance (Delorme and Makeig, 2004; Antonakakis et al., 2013). The resulting signals were further submitted to ICA using the extended Infomax algorithm as implemented in EEGLAB (Delorme and Makeig, 2004). A given independent component (IC) was considered to reflect ocular,muscle or cardiac artifacts if more than 30% of its z-score kurtosis or skewness values, respectively, were outside ±2 of the distribution mean (Antonakakis et al., 2013; Dimitriadis et al., 2015c). Finally, the artifactual IC were zeroed and the artefact free activity was back-projected to the original dimension space. ICA was used as the candidate denoising method for every dataset.

### 2.2.2 EEG Analysis

EEG activity of {δ, θ, α_1_, α_2_, β_1_, β_2_, γ} frequency bands defined respectively within the ranges {0.5–4 Hz; 4–8 Hz; 8–10 Hz; 10–13 Hz; 13–20 Hz; 20–30 Hz; 30–48 Hz}. EEG recordings were bandpass filtered with a 3^rd^ order zero-phase Butterworth filter using filtfilt.m MATLAB function.

### 2.2.3 ECoG Analysis

ECoG activity of {δ, θ, α_1_, α_2_, β_1_, β_2_, γ_1_, γ_2_ } frequency bands defined respectively within the ranges {0.5–4 Hz; 4–8 Hz; 8–10 Hz; 10–13 Hz; 13–20 Hz; 20–30 Hz; 30–48 Hz }. ECoG recordings were bandpass filtered with a 3^rd^ order zero-phase Butterworth filter using filtfilt.m MATLAB function.

### 2.2.4 Beamformer Analysis of MEG Activity

The activity of {δ, θ, α_1_, α_2_, β_1_, β_2_, γ } frequency bands defined respectively within the ranges {0.5–4 Hz; 4–8 Hz; 8–10 Hz; 10–13 Hz; 13–20 Hz; 20–30 Hz; 30–48 Hz } was first beamformed with linear constrained minimum norm variance (LCMV) in the artefact-free MEG data to determine ninety anatomical regions of interest (ROIs) in a template volume conduction head model. Here, we used the AAL-90 ROIs atlas.

The beamformer sequentially reconstructs the activity for each voxel in a predefined grid covering the entire brain (spacing 6 mm) by weighting the contribution of each MEG sensor to a voxel’s time series a procedure that creates the spatial filters that can then project sensor activity to the cortical activity. Every ROI within the cortex contains many voxels and there are many algorithms of how to represent each ROI with a representative time series. Here, we estimated the representative time series via a linear weighted interpolation of the entire set of voxel time series that are encapsulated within every ROI. This method has already been demonstrated in a previous study employing the same MEG dataset (Dimitriadis et al., 2018).

The whole analysis was written in MATLAB with routines from fieldtrip toolbox (Oostenveld et al., 2011). For further details regarding MRI acquisition and beamforming analysis, see (Dimitriadis et al., 2018).

### 2.2.5 fMRI Recordings

Artifact free Fmri recordings were decomposed using the maximal overlap discrete wavelet transform (MODWT) method to the related wavelet coefficients for the first four wavelet scales, which in this case correspond to the frequency ranges 0.125~0.25 Hz (Scale 1), 0.06~0.125 Hz (Scale 2), 0.03~0.06 Hz (Scale 3), and 0.015~0.03 Hz (Scale 4) (Dimitriadis et al., 2017). Bold activity of each of the 630 regions was decomposed with MODWT in wavelet coefficients for each scale. Free-surfer parcellation of BOLD activity gave a total of 630 regions for subsequent analysis. For further details please see the original paper (Poldrack et al., 2015).

### 2.3 Neural-Gas algorithm and Complexity Index (CI)

An alternative method to transform the time-series expressed the brain activity into symbols is to adopt a proper algorithm that can learn the manifold of a reconstructed phase space and then determining the appropriate mapping between trajectories and symbols (alphabet). Here, i embedded each band-passed time-series (Fig.3.A.) into a common reconstructed space (Fig.3.B) and then applied the NG algorithm to derive a set of symbols that can describe the original signal with a reduced amount of error (Fig.3.C). For details on the procedure see (Dimitriadis et al., 2016d). Each concatenated time series was first embedded in a multidimensional space as described in equation (1):

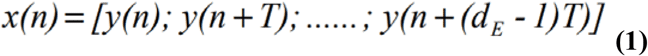

where the time lag T is determined using mutual information and the embedding dimension dE is obtained using the false nearest neighbors test (Abarbanel, 1996). Having estimated the reconstructed error between the original MEG time series and the one described by the NG-derived codebook, we fixed the number of symbols for each time series. The reconstruction error was set equals to 0.04. Finally, each ROI-based time-series was transformed to a Symbolic Time Series STSNG=[1 2 3 4 5 6 2 1 …] where each integer corresponds to one symbol (Fig.3.A-D).

**Figure 3.**
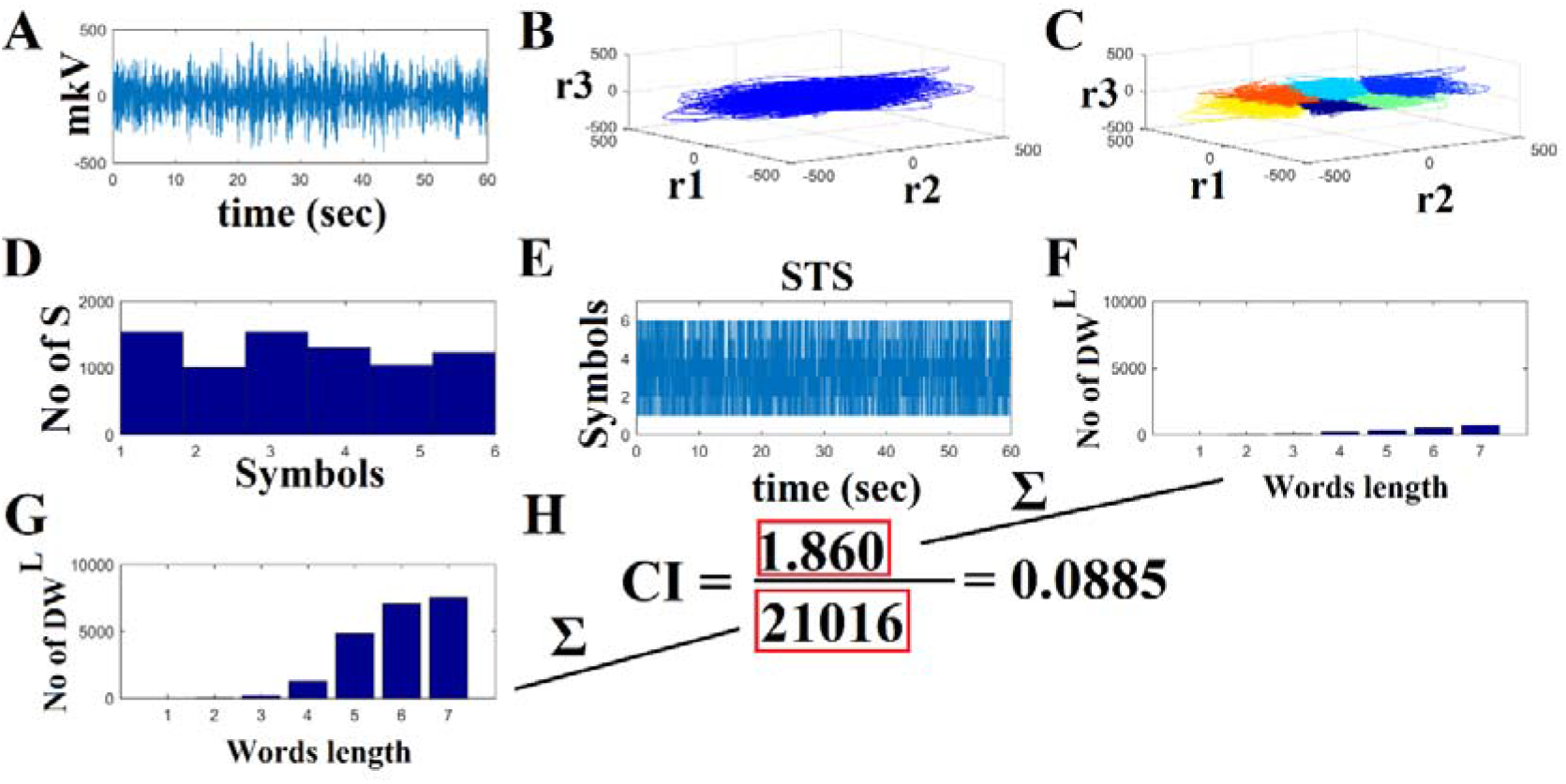
From raw time-series to CI. A. The original EEG activity in θ band from the first healthy control subject over Fp1 sensor B. The reconstructed embedded space of the time series demonstrated in A using embedding dimension 3 and time-delay equals to 9. C. Applying neural-gas algorithm in B, we clustered the time-points in the embedding space into 6 classes where each corresponds to a symbol. D. The distribution of the Symbols S across the embedding space. E. Symbolic Time Series (STS) as the outcome of the neural-gas algorithm demonstrated in C F. Distribution of distinct words length up to length 7 for the real STS demonstrated in E G. Mean distribution of distinct words up to length 7 from 1000 randomized versions of the original STS H. CI is estimated as the ration of the sum of distribution of distinct words up to length 7 illustrated in F versus the mean distribution of distinct words up to length 7 from the 1000 randomized versions of the original STS as shown in G.

We applied this learning scheme within each group and frequency independently of each ROI activity for EEG,MEG and fMRI datasets.

CI quantifies the “richness of the language” within a symbolic sequence, and has been used in several fields, such as data compression, data mining, computational biology computational linguistics (Leve and Seebold, 2001). We adopted the CI based on symbolic sequences as presented in Janson et al. (2004). Low CI values describe sequences containing frequent repeated substrings that become periodic.

The magnitude of the derived CI values was normalized based on the deviation from the complexity that can be derived by random-shuffled versions of the original symbolic sequence. Here, we used 1,000 randomized versions of the original symbolic sequence and the normalization procedure can be seen in Fig.3.G (Dimitriadis et al., 2016b).

### 2.4 Dynamic iPLV estimates: the time-varying integrated iPLV graph (^TVI^iPLV graph)

The goal of the analytic procedures described in this section is to understand the repertoire of phase-to-phase interactions and their temporal evolution, while taking into account the quasi-instantaneous spatiotemporal distribution of iPLV estimates. This was achieved by computing one set of iPLV estimates within each of a series of sliding overlapping windows spanning the entire recording set for continuous EEG-ECoG-MEG recording for the three out of four datasets. For fMRI dataset, I adopted wavelet coherence (WC) estimated over the wavelet decomposed time series in order to avoid positive and negative values of the correlation.

The width of the temporal window was set equal to the duration of 1 sec, as an adequate number to capture the dynamics of every brain rhythm (fast and slows, Dimitriadis et al., 2013a). For EEG/ECoG/MEG, the center of the stepping window moved forward every 20 ms and both intra and inter-frequency interactions between every possible pair of frequencies were re-estimated leading to a quasi-stable in time static iPLV graph. For fMRI dataset, i adopted a time-window of 20 TR and the center of window moved forward by step equals to 1 TR and in every temporal segment, quasi-static WC graph were estimated that tabulates functional interactions within and between frequencies between every possible pair of ROIs.

In this manner, a series of iPLV graph estimates were computed per subject or condition, for each of the intra-frequency coupling (7 (EEG) or 8 (ECoG-MEG) or 4 (fMRI)-frequencies) and the 21 (EEG) or 28 (ECoG-MEG) or 6 (fMRI) possible cross-frequency pairs.

This procedure, the implementation details of which can be found elsewhere (Dimitriadis et al., 2010, 2012, 2013b, 2015a,c), resulted in 7 (EEG) or 8 (ECoG-MEG) or 4 (fMRI) time-varying graphs per participant (^TV^iPLV or ^TV^WC) for within frequency bands and 21 (EEG) or 28 (ECoG-MEG) or 6 (fMRI)) ^TV^iPLV or ^TV^WC graphs per participant for each possible cross-frequency pair, each serving as an instantaneous snapshot of the surface network. ^TV^iPLV or ^TV^WC tabulate iPLV/WC estimates between every possible pair of sensors/sources/ROIs. For each subject, a 4D tensor [frequencies bands (28 (EEG),36(ECoG-MEG), 10 (fMRI) × slides × sensors/sources/ROIs × sensors/sources/ROIs] was created for each condition integrating subject-specific spatio-temporal phase interactions.

Afterward, we applied surrogate analysis in order to reveal the dominant type of interaction for each pair of sensors/sources/ROIs and at each snapshot of the ^TV^iPLV or ^TV^WC. We constructed 1000 surrogate time-series by cutting first at a single point at a random location the original time series, creating two temporal segments and then exchanging the two resulting temporal segments (Aru *et al.,* 2015). We restricted the range of the selected cutting point in a temporal window within the middle of the recording session. This procedure alters the temporal coherence between the pairs of every temporal segment for every possible coupling mode. The proposed surrogate scheme was applied to the original whole time series and not to the signal-segment at every slide. Repeating this procedure leads to a set of surrogates with a minimal distortion of the original phase dynamics (see Dimitriadis et al., 2017). Finally, for each pair of sensors/sources/ROIs and for each temporal segment, I estimated 1000 iPLV/WC for every within frequency interaction and every possible pair of frequencies. Practically, we assigned a p-value to each within and between frequencies interaction and for each sensors/sources/ROIs by comparing the original iPLV/WC value with 1000 surrogates iPLV^Surr^/WC^iPLV^. Then, we corrected for multiple comparisons across 28 (EEG),36(ECoG-MEG) and 10 (fMRI) possible DICM in order to reveal a DICM per pair of sensors/sources/ROIs and for each temporal segment. There are in total three scenarios:

a. no p-value survived the multiple correction (p’ < p/(28 or 36 or 10) where p=0.05)
b. more than one survived and in that case, we selected the DICM with the maximum iPLV/WC value or
c. only one survived the multiple correction

Fig. 4. A illustrates how the DICM is defined in the first two temporal segments from the EEG dataset between FP1 and P4 EEG sensors.

**Figure 4.**
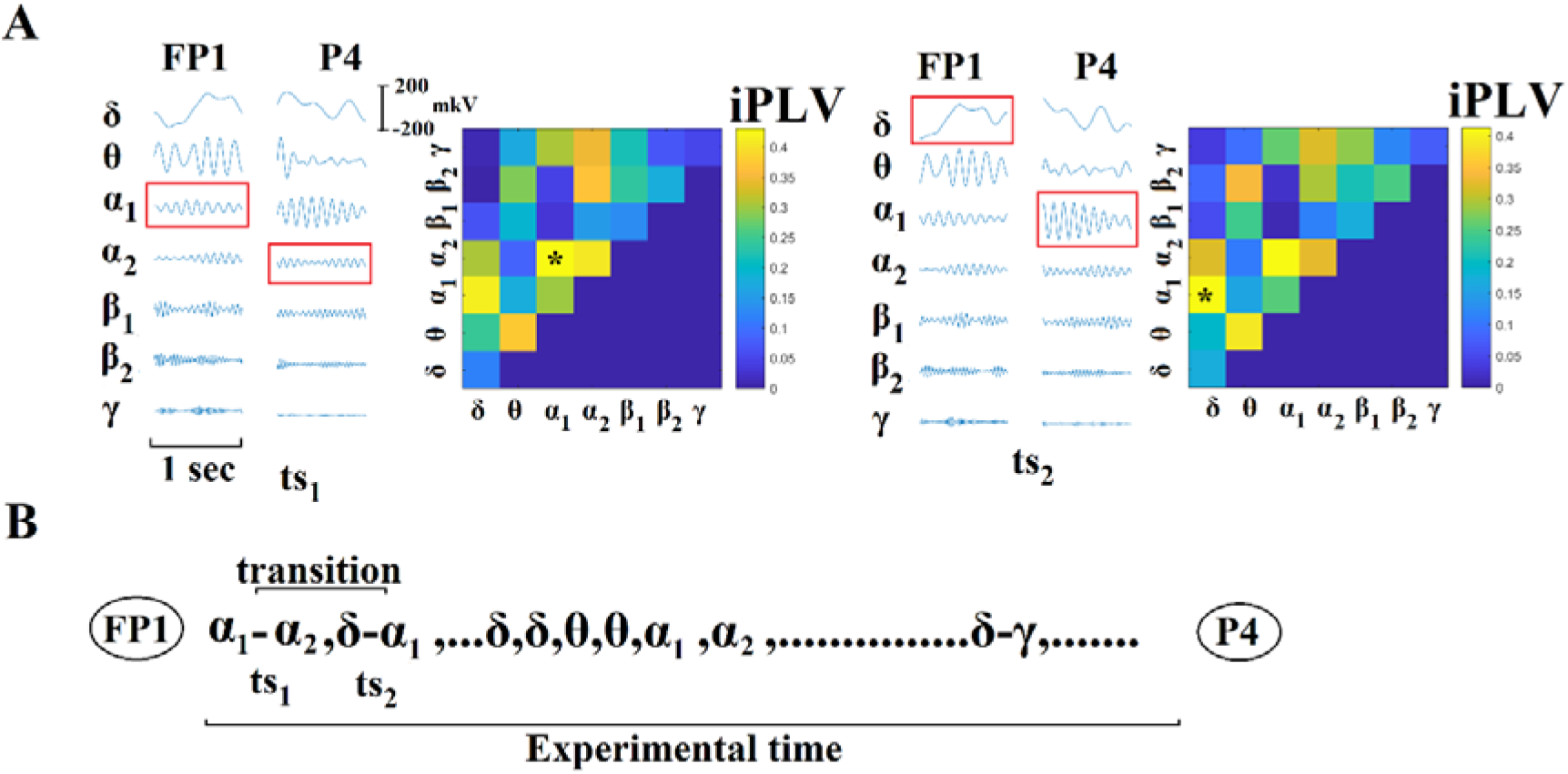
Determining Dominant Intrinsic Coupling Modes (DICM). A. Schematic illustration of the approach employed to identify the DICM between two EEG sensors (FP1 and P4) for two consecutive 1s sliding temporal segment (ts_1_, ts_2_) during the resting-state EEG activity from the first normal subject. In this example the functional interdependence between band-passed signals from the two sensors was indexed by imaginary Phase Locking (iPLV). In this manner iPLV was computed between the two EEG sensors either at same-frequency oscillations (e.g., δ to δ) or between different frequencies (e.g., δ to θ). Statistical filtering, using surrogate data for reference, was employed to assess whether each iPLV value was significantly different than chance. During ts_1_ the DICM reflected significant phase locking between α_1_ and α_2_ oscillations (indicated by red rectangles) whereas during in ts2 the dominant interaction was found between δ and α_1_ oscillations. B. Burst of DICM between the two sensors. This packeting can be thought to associate the ‘letters’ contained in the DICM series to form a neural “word.”, a possible integration of many DICM (Buzsaki and Watscon,2012). For the first pair of ts1-2, i illustrated how a transition is defined for FP1-P4 EEG pair important for the estimation of FI (see section 2.5).

Finally, we tabulated both the strength and the type of dominant coupling mode in 2 3D tensor [slides × sensors/sources/ROIs × sensors/sources/ROIs], one that keeps the strength of the coupling based on iPLV/WC and a second one for keeping the dominant coupling of interaction using integers from 1 up to 28 (EEG),36 (ECoG-MEG) or 10 (fMRI) {for ECoG-MEG: 1 for δ, 2 for θ,…,8 for γ_2_, 9 for δ–θ,…, 36 for β_2_ – γ_2_, for fMRI: 1 for scale 1, 2 for scale 2, …, 10 for scale 3-scale 4 }. The notion of phase-to-amplitude cross-frequency coupling (CFC) estimator has been used also in our previous studies with EEG/MEG brain signals (Dimitriadis et al., 2015b, 2016a,b,c, 2017).

### 2.5 Dominant intrinsic coupling mode transition rate

Based on the 2nd 3D DIFCG that keeps the information of the DICM per pair of sensors/sources/ROIs and across time, we estimated the transition rate for each pair of sensors/sources/ROIs. The estimator is called flexibility index (FI) proposed and quantifies how many times a sensor/source/ROI changes DICM across experimental time similar but not the same with Flexibility Index based on cluster assignment (Bassett et al., 2011). This metric will called hereafter FI^DICM^ which is defined as:

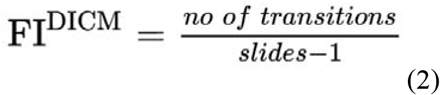

FI^DICM^ gets higher values for higher “jumps” of DICM between a pair of sensors/sources/ROIs between consecutive time windows. Fig.4.B illustrates how a transition is estimated for the first pair of temporal segments ts_1-2_ and for FP1-P4 EEG pair.

This approach leads to [sensors/sources/ROIs] x sensors/sources/ROIs features per subject or scan.

### 2.6 Spatio-temporal distribution of dominant intrinsic coupling modes— comodulograms

Based on the 2^nd^ DIFCG that keeps the information of the DICM, we can tabulate in a frequencies × frequencies matrix the probability distribution of observing each of the DICM frequencies (7 (EEG) or 8 (ECoG-MEG) or 4 (fMRI)–frequencies + 21 (EEG) or 28 (ECoG-MEG) or 6 (fMRI)-cross-frequency pairs) across space and time.

This spatio-temporal probability distribution is called hereafter as comodulograms. For further details see Dimitriadis et al., 2016a,2017.

### 2.7 Reliability of FI

The reliability of FI has been accessed with intra-class correlation (ICC) index (Koo and Mae, 2016).

## 3. Results

In this section, we reported the results of CI,FI and comodulograms for each functional neuroimaging dataset separately.

### 3.1 Classifying Healthy Controls and Schizophrenic Adults with CI and FI

We applied a feature selection and machine learning strategy independently for CI, FI and comodulograms. We employed the laplacian score as a feature selection algorithm (He et al., 2005) and kNN classifier with 5 nearest neighbors as a simple classifier in order to further enhance the discriminative power of the novel complexity indexes. We used a 5-fold crossvalidation scheme using the 75% of subjects from both groups in order to internally optimize the number of features. kNN classifier was trained in the 75% of the dataset and tested on the rest 25% with pre-selected number of features.

Fig. 5 illustrates the group-averaged CI across EEG sensors and frequency bands. The selected seven CI features are denoted with ‘*’. The classification performance based on CI reached 78%.

**Figure 5.**
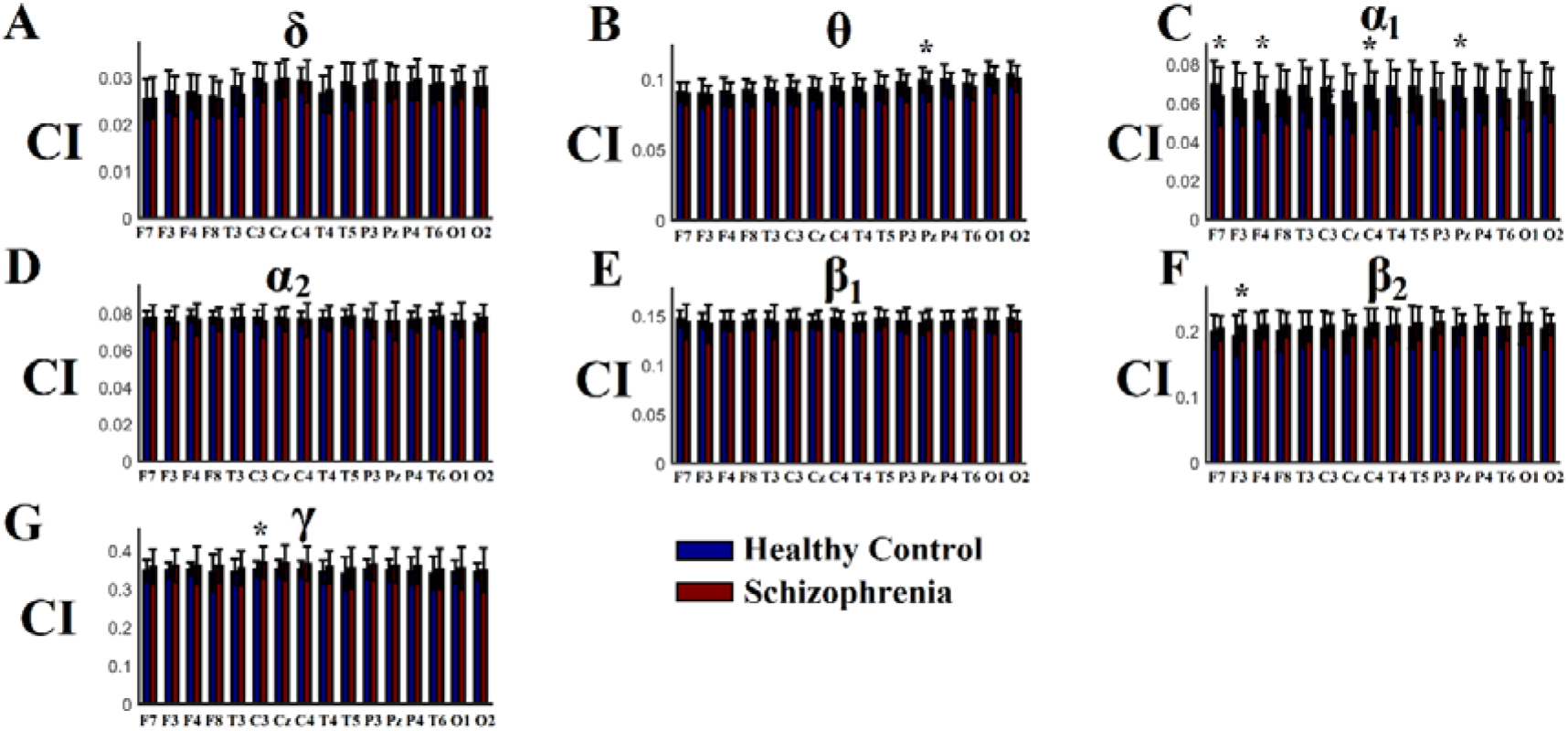
Group-Averaged Complexity Index (CI) across EEG sensors. A-G) Group-averaged CI for δ up to γ frequency. Every CI selected is denoted with ‘*’.

Fig. 6.A-B illustrates the group-averaged FI across every pair of EEG sensors for healthy control and schizophrenic group,correspondingly. The selected ten FI features (connections) are denoted with ‘*’ and are located in fronto-temporo-parietal network. The classification performance based on those ten FI was absolute (100%).

**Figure 6.**
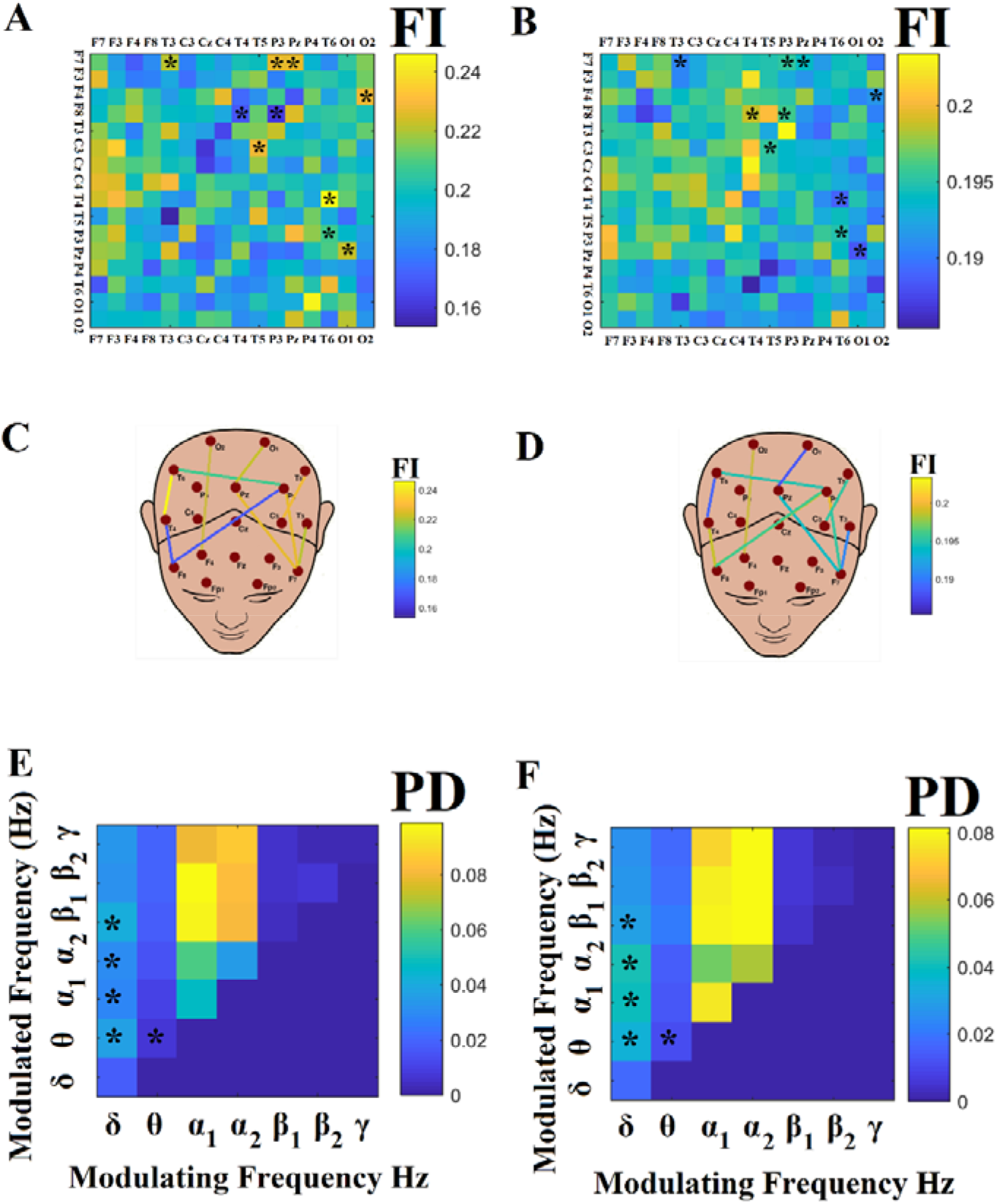
Group-Averaged Flexibility Index (FI) and Comodulograms. A-B) Group-averaged FI for healthy control group (A) and schizophrenic group (B) C-D) Group-averaged comodulograms for healthy control group (C) and schizophrenic group (D) estimated across space and time. Every FI and PD selected is denoted with ‘*’.

Fig. 6.C-D illustrates the group-averaged comodulograms for healthy control and schizophrenic group, correspondingly. The selected five comodulograms features are denoted with ‘*’ and are referred to the probability distribution (PD) across space and time of δ-θ, δ-α_1_,δ-α_2_, δ-β_1_ and θ-α_1_. The classification performance based on selected PD features was absolute (100%).

### 3.2 Sensitivity of FI and Comodulograms of DICM during Awake and Anesthesia

Fig. 7 illustrates the group-averaged CI for each frequency band and for both awake and anesthesia conditions. CI values were first averaged across the ECoG sensors for each condition and monkey and afterward across the cohort. Statistical significant trends were detected between the two conditions (p < 0.01, Wilcoxon rank sum test, Bonferroni corrected, p’<p/8). CI was higher in awake condition compared to anesthesia condition in all frequency bands with the exception of δ while CI was higher in anesthesia compared to awake condition in δ frequency.

**Figure 7.**
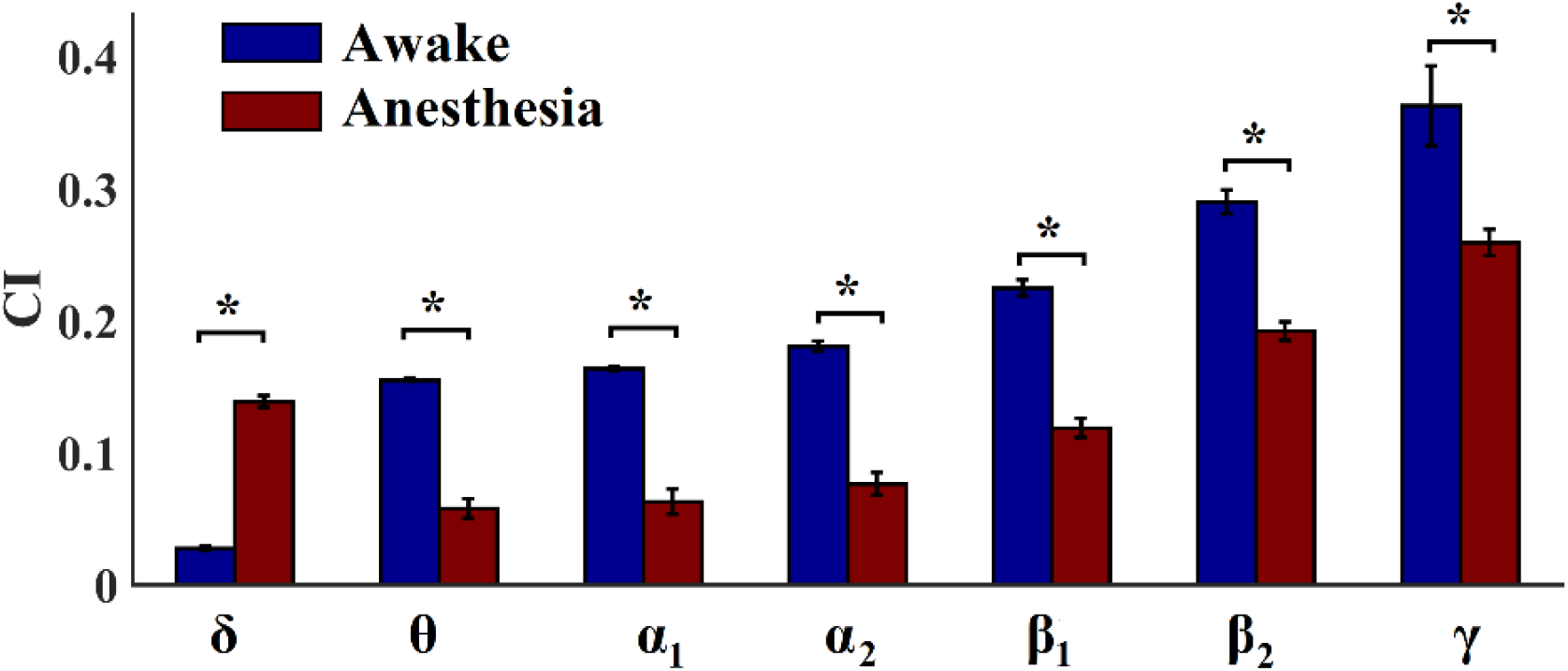
Group-Averaged Complexity Index (CI) across ECoG sensors and monkeys. A-G) Group-averaged CI for up to frequency. Every CI that statistically differed between the two conditions is denoted with ‘*’. (p < 0.01, Wilcoxon rank sum test, Bonferroni corrected, p’<p/8)

Fig. 8.A-B illustrates the group-averaged topologies of FI across every pair of ECoG sensors for awake and anesthesia group,correspondingly. We applied a z-score > 3 to every condition in order to enhance the visualization of the survived connections. It is clear that FI is reduced during anesthesia while a dense network is located over higher and lower visual areas with a few connections between visual areas and lateral prefrontal cortex during anesthesia (Fig.8.A) compared to a dense network during awake condition (Fig.8.A).

**Figure 8.**
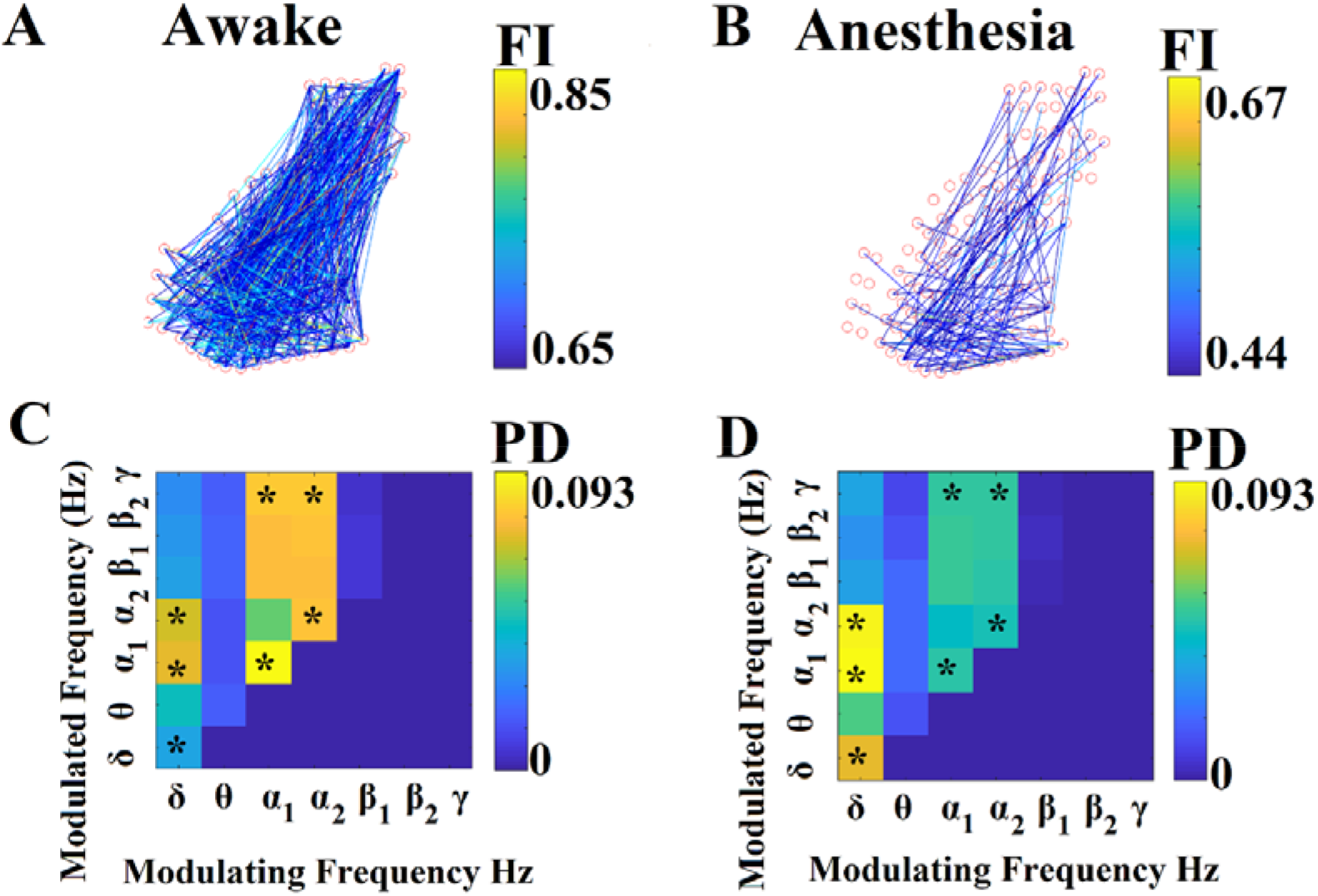
Group-Averaged Flexibility Index (FI) and Comodulograms. A-B) Group-averaged topologies FI for awake idle condition (A) and for anesthesia (B). We applied a z-score > 3 as a threshold to enhance the visualization of the survived connections based on FI. C-D) Group-averaged comodulograms for awake condition (C) and anesthesia (D) estimated across space and time. Every FI and PD selected is denoted with ‘*’.

The selected twelve FI features (connections) via the machine learning scheme are located between medial prefrontal cortex, later prefrontal cortex and parietal cortexThe classification performance for separating awake from anesthesia condition based on FI was absolute (100%).

Fig. 8.C-D illustrates the group-averaged comodulograms for awake and anesthesia conditions, correspondingly. The selected seven comodulograms features are denoted with ‘*’ and are referred to the probability distribution (PD) across space and time of δ-α_1_,δ-α_2_ (higher for anesthesia), α_1_, α_2_, α_1_-γ, α_2_-γ (higher in awake) and δ (higher in anesthesia). The classification performance between awake and anesthesia based on PD was absolute (100%).

### 3.3 Reliability of FI and Comodulograms of DICM during MEG Resting-State

Figure 9 illustrates the group-averaged mean CI values across the frequency bands and within the five well-known brain networks. CI values were first averaged within each brain network and for each scan session afterward across scans and the standard deviation was estimated across subjects. Additionally, intra-class correlation values have been estimated in order to access the reproducibility of CI values (Fig.9.B).

**Figure 9.**
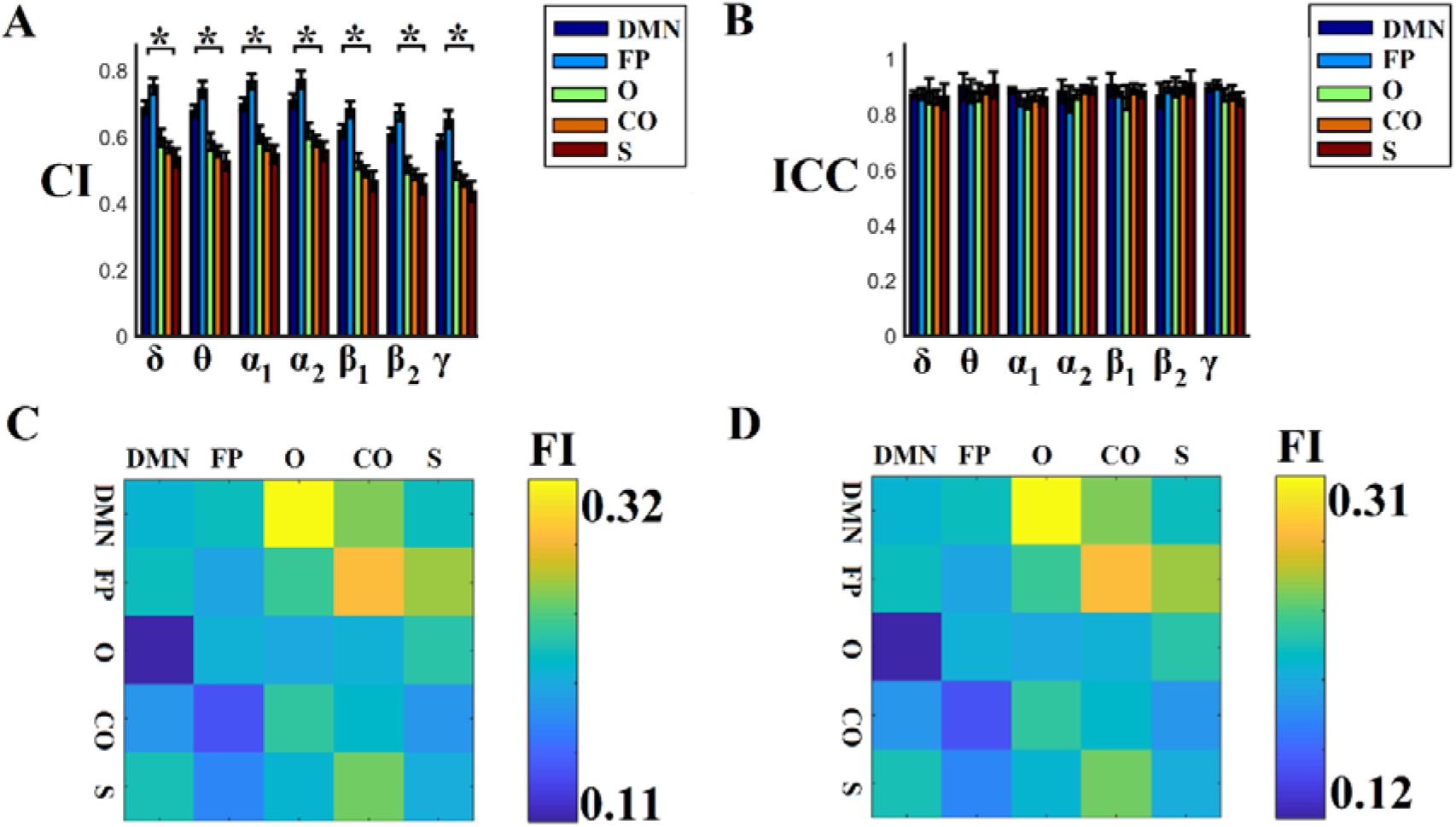
Group-Averaged Complexity Index (CI) and Flexibility Index (FI) Group-averaged CI for δ up to γ_2_ frequency. A) Group-averaged CI for δ up to γ frequency. B) Group-averaged ICC for δ up to γ frequency. C) Group-averaged FI within and between brain networks from the first scan session D) Group-averaged FI within and between brain networks from the second scan session

The adopted tatistical analysis revealed a common frequency-dependent trend where DMN and FP demonstrate significant higher CI compared to the rest of brain networks across the frequency bands (Fig.9.A, p < 0.01, Bonferroni corrected p’<p/80 where 80 denotes the 8 frequency bands multiplied by the ten pair-wise comparisons between every pair of brain networks). Complementary, the CI was higher for the lower frequency bands (δ,θ,α1,α2) compared to the higher (β_1_, β_2_, γ).

Finally, CI values were high reliable across the frequency bands and brain networks with ICC values above 0.85 (Fig.9.B).

FI values estimated within and between brain networks were also high reliable between scan session 1 and scan session 2 (Fig.9. C,D). Interestingly, the more flexible set of pairs of brain networks are the DMN-CO, FP-CO following by DMN-CO and FP-S.

Every CI that statistically differed between the two conditions is denoted with ‘*’. (p < 0.01, Bonferroni corrected p’<p/80 where 80 denotes the 8 frequency bands multiplied by the ten pair-wise comparisons between every pair of brain networks). (DMN:Default Mode Network, FP:Fronto Parietal, O:Occipital, CO:Cingulo-Opercular, S:Sensorimotor)

In Fig.10.A,B the group-averaged comodulograms are illustrated for scan session 1 and 2, correspondingly. Similarly, in Fig. 11.C,D, the group-averaged comodulograms are demonstrated within and between every pair of brain networks for scan session 1 and 2, correspondingly. It is worth to notice that the pattern of DICM is highly reproducible in both spatial scales (whole network A-B and between networks C-D). Our methodology harness the notion of DICM in order to reveal the multiplexity of human brain dynamics in both spatial scales using MEG resting-state activity. Δ and α2 frequencies govern the DICM and the multiplexity of neuromagnetic recordings at resting-state. The estimated ICC for both estimated comodulograms (Fig.10) was 0.91 ± 0.06 across the cohort.

**Figure 10.**
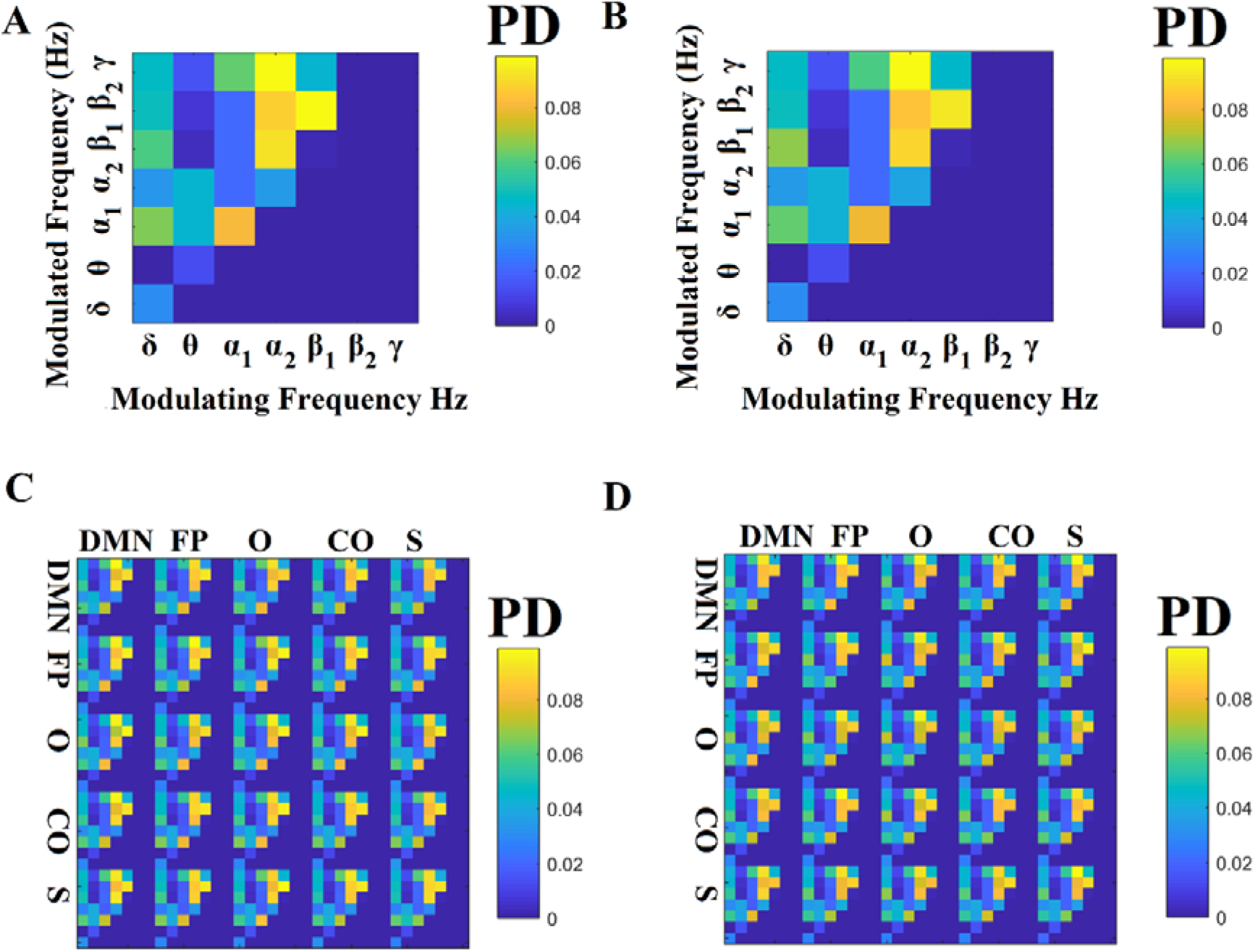
Group-Averaged Comodulograms in whole brain and brain networks. A-B) Group-averaged comodulograms for scan session 1 (A) and scan session 2 (B) C-D) Group-averaged comodulograms within and between the five brain networks for scan session 1 (C) and scan session 2 (D). (DMN:Default Mode Network, FP:Fronto Parietal, O:Occipital, CO:Cingulo-Opercular, S:Sensorimotor)

**Figure 11.**
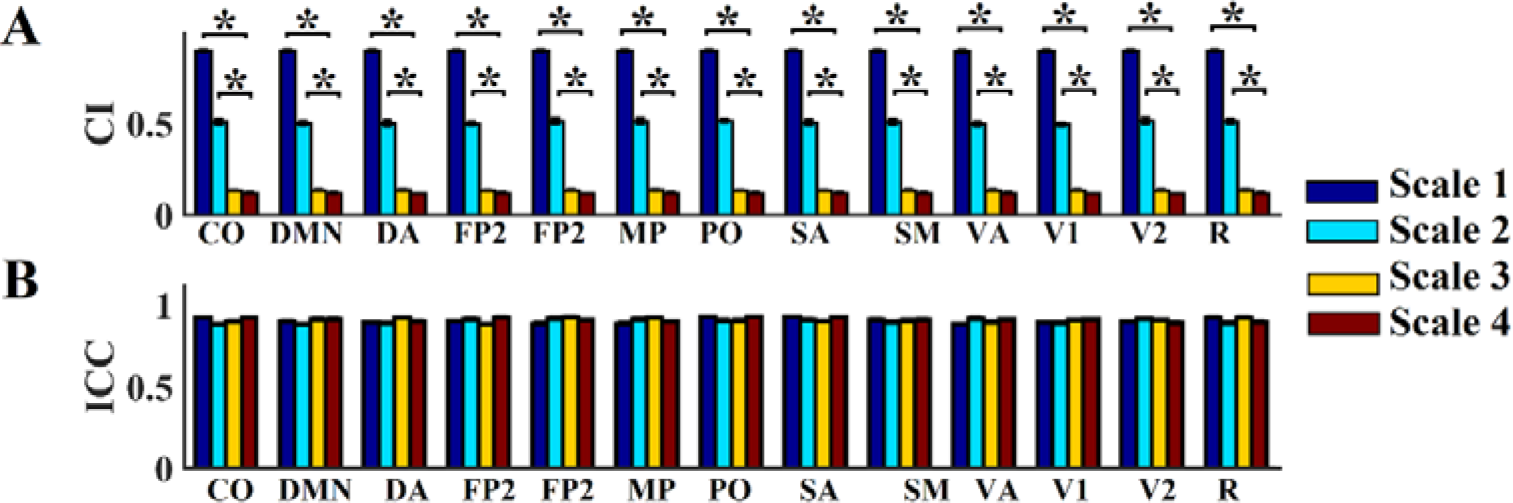
Scan-Averaged Complexity Index (CI) and ICC. A) Scan-Averaged Complexity Index (CI) B) ICC values across frequency sub-bands (scales) and the thirteen brain networks Statistical significant differences are denoted with ‘*’ (*, p < 0.01, Bonferroni corrected p’<p/78 where 78 denotes the thirteen brain networks multiplied by the six pair-wise comparisons between every pair of brain networks). CO:Cingulo-opercular, DMN:Default Mode Network, DA: Dorsal Attention, (FP1:Frontoparietal 1, FP2:Frontoparietal 2, MP:Medial-Parietal, PO:Parieto-Occipital, S:Salience, SM:Somatomotor, VA: Ventral-Attention,V1:Visual-1, V2:Visual-2, R:rest of unclassified nodes)(p < 0.01, Bonferroni corrected p’<p/8 where 78 denotes the thirteen brain networks multiplied by the six pair-wise comparisons between every pair of brain networks).

### 3.4 Reliability of FI and Comodulograms of DICM during fMRI Resting-State

Fig.11.A illustrates the scan-averaged CI estimated within thirteen brain networks and the related ICC for fMRI resting-state. CI was statistically significant different between scale 1 and the rest 2-4 and also between scale 2 and scales 3-4 across brain networks (p < 0.01, Bonferroni corrected p’<p/78 where 78 denotes the thirteen brain networks multiplied by the six pair-wise comparisons between every pair of brain networks). It seems that CI follows the frequency scales with the exception for scales 3 which had similar CI values with scale 4. ICC values were > 0.9 meaning that CI were high reliable (Fig. 11.B).

Fig.12.A and B demonstrates the scan-averaged FI between and within the thirteen brain networks in both split-half scan sessions. The outcome clearly supports the reliability of the FI which was *ICC =* 0.93 ± 0.03 across scan – sessions.

Trial-averaged comodulogram across the whole brain network and between and within the thirteen brain networks are illustrated in Fig.12.C and D, correspondingly. Fig.12.C reveals frequency scale 1 as the basic brain modulator following by scale 2. These pattern was observed also on the more detailed spatial scale of brain networks both within (main diagonal) and between brain networks (off-diagonal) (Fig.12.D).

**Figure 12.**
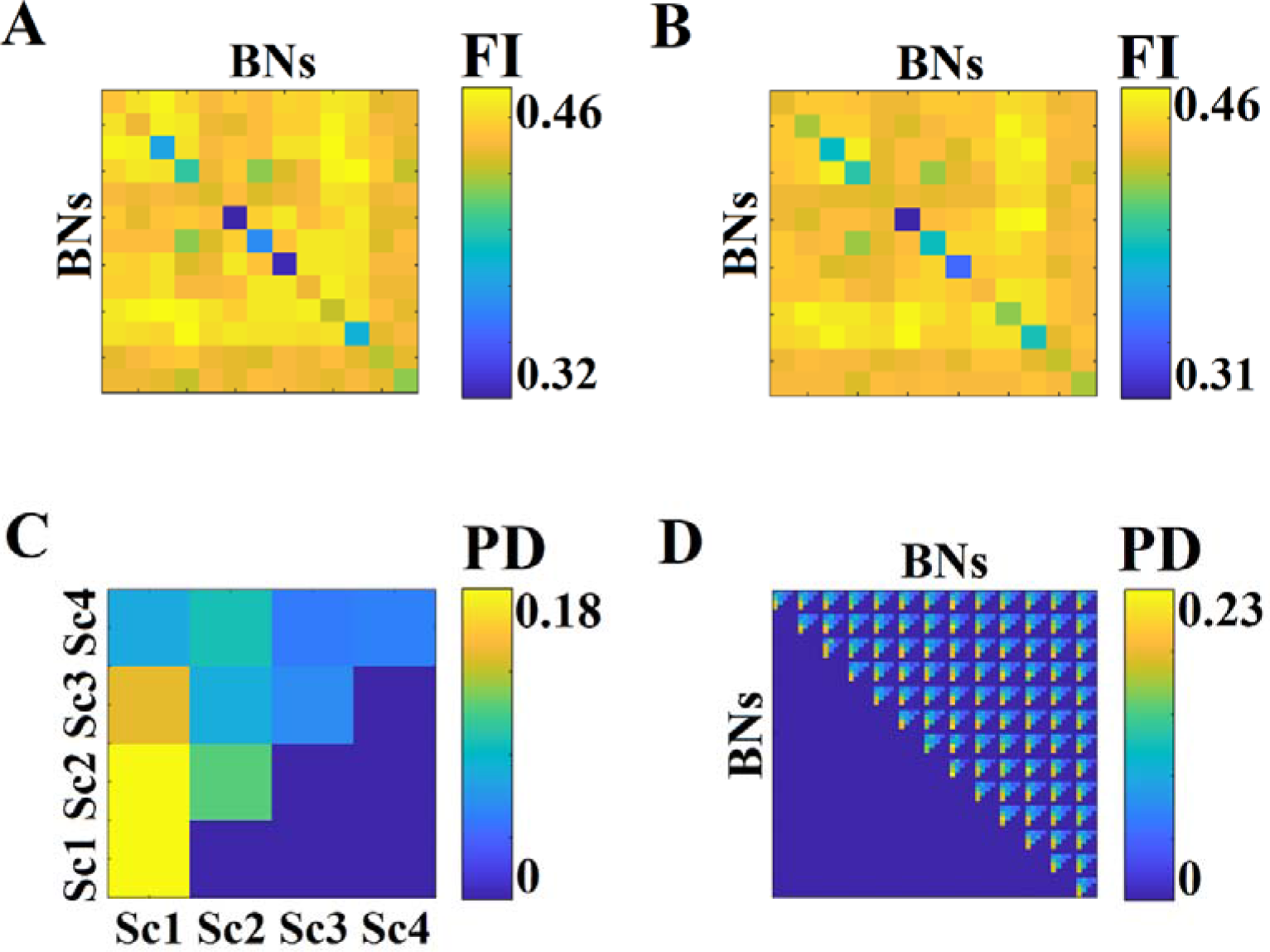
FI and Comodulograms across the repeat scans fMRI resting-state. A. FI within and between the brain networks for the first half of repeat scans B. FI within and between the brain networks for the second half of repeat scans C. Scan-averaged comodulograms estimated across the whole network D. Scan-averaged comodulograms estimated within and between the brain networks

## 4. Discussion

In the present study, we presented a methodology approach for the estimation of complexity of brain activity and also the complexity of brain connectivity under the notion of dominant intrinsic coupling modes (DICM). We decided to demonstrate the whole approach using representative datasets from the majority of functional neuroimaging modalities. As a representative EEG dataset, we selected one with healthy controls and schizophrenic patients at resting-state in order to demonstrate the effectiveness of the proposed methodology to discriminate the two populations. ECoG recordings were selected from four monkeys undergoing anesthesia with different drugs in order to reveal the different pattern of multiplexity and DICM between awake and anesthesia and also a common trend independently of the drug. Reliability is very important nowadays for functional neuroimaging. For that reason, we have analysed repeat-MEG and fMRI scan recordings in order to estimated the reliability of CI,FI and DICM across the repeat scans.

There are several complexity indexes that can quantify the randomness of sequence. The most famous is called LempelZiv complexity of a sequence which was defined by Lempel and Ziv and it is known as LZ complexity index (Lempel and Ziv, 1976). This complexity measure counts the number of different patterns in a symbolic binary sequence with a finite length when scanned from the left to right. For example, the Lempel-Ziv complexity of the sequence s = 101001010010111 is 7, equals the total number of different patterns 1|0|10|01|010|0101|11| when scanned from the left to the right. The disadvantage of this algorithm is the arbitrary selection of the threshold that is needed for the binarization of the original time series into a symbolic sequence of 0s and 1s. Moreover, LZ complexity index cannot reveal the complex of a non-linear system described by a time series. For the aforementioned reasons, we introduced here a novel approach which first embedded the time series into a reconstructing state space and then applying neural-gas algorithm, we clustered the data time points into a specific group of points selected in a data-driven way via the reconstruction error. Based on the derived symbolic sequence, we estimated the spectrum of the total number of words up to a length. The same spectrum is estimated for a number of surrogated symbolic time series produced by shuffling the original time series. The novel complexity index is estimated by dividing the original spectrum with the sum of spectrums estimated from the surrogates (Fig.3). In two previous studies, we tested the proposed CI versus the LZ complexity index. We reported an improved classification accuracy with the proposed CI compared to LZ index of dyslexic children versus non-impaired readers (Dimitriadis et al., 2017) and of mild traumatic brainn injured patients versus healthy controls using (Antonakakis et al., 2017). We also found out higher CI values for dyslexic children versus non-impaired readers using MEG resting-state (Dimitriadis et al., 2017) and lower CI values for mTBI subjects versus healthy controls using MEG resting-state (Antonakakis et al., 2017).

We designed a novel flexibility index (FI) based on the spatio-temporal fluctuation of dominant intrinsic coupling modes (DICM) in order to explore the multiplexity of human brain dynamics. Till now, brain connectivity has been studied independently for each brain rhythm avoiding also to explore cross-frequency interactions across the whole brain. Here, we introduced to the neuroscience community, a way of how to integrate into a single dynamic integrated functional connectivity graph the repertoire of possible coupling modes. This repertoire includes both intra and cross-frequency coupling mechanisms. Human brain mechanisms support distinct temporal frames to group brain activity into sequences of accemblies where multi-frequency interactions occur across the whole brain. These multiplex interactions create the syntactic rules which are significant for the exchange of information across the cortex (Buzsaki and Watscon,2012). Here, we provided a framework of how to study the majority of available and well established interactions simultaneously and across the whole brain. It is important to explore brain interactions globally and afterward to focus on sub-networks and local interactions. On the top of it, we define an index that quantifies the flexibility of a pair of ROIs which quantifies the exchange rate of the preferred (dominant) coupling mode.

Both measures of CI and FI have been presented in the majority of neuroimaging studies while they can be adapted in any study at resting-state and also on task-related activity with any neuroimaging modality. Moreover, multi-modal correlation of CI – FI values derived from datasets acquired from simultaneously recordings of two functional modalities like in EEG-MEG, EEG-fMRI is more than significant. It would more than important to explore the enriched information of both indexes to build a sensitive biomarker for a large number of brain disorders/diseases like in dyslexia (Dimitriadis et al.,2017), in mild traumatic brain injury (Antonakakis et al., 2017), in Alzheimer’s disease, in schizophrenia etc.

Following a machine learning approach, we revealed very interesting results related to the novel introduced features that can potentially discriminate the healthy controls from the schizophrenia patients. CI values succeeded to classify the two groups with 78% while FI and comodulograms with absolute accuracy (100%). The selected FI features were topologically located between fronto-temporo-parietal brain areas. Schizophrenia alters the probability distributions (PD) of δ-θ, δ-α_1_,δ-α_2_ δ-β_1_ and θ-α_1_. A recent study revealed frontal slow-wave abnormalities in schizophrenia that are associated with negative symptoms while the increase of frontal δ activity in schizophrenic populations is linked to poorer attention (Chen et al., 2015). Complementary, schizophrenia alters the dynamic reconfiguration of DICM which is reflected in both FI and comodulograms. This is the very first study that reports significant results in schizophrenia under the notion of multiplexity including cross-frequency coupling estimates. It’s more than important to gain the advantage of functional neuroimaging and especially the EEG/MEG to reveal the dominant coupling modes in schizophrenia and in general in psychiatry (Alamian et al., 2017).

The analysis of ECoG recordings from four monkeys in awake condition and during anesthesia with various drugs untangled significant common trends. First of all, CI was higher in awake condition compared to anesthesia condition in all frequency bands with the exception of δ where CI was higher in anesthesia compared to awake condition (Murphy et al., 2011).

Our analysis revealed a reduced FI during anesthesia while a dense network is located over higher and lower visual areas with a few connections between visual areas and lateral prefrontal cortex compared to the more dense network during awake.

Machine learning approach selected twelve FI features (connections) which are located between medial prefrontal cortex, later prefrontal cortex and parietal cortex. The classification performance for separating awake from anesthesic condition based on FI was absolute (100%).

Additionally, machine learning scheme selected seven comodulograms features referred to the probability distribution (PD) across space and time of δ-α_1_,δ-α_2_ (higher for anesthesia), α_1_, α_2_, α_1_-γ, α_2_-γ (higher in awake) and δ (higher in anesthesia). The classification performance between awake and anesthesic conditions based on PD was absolute (100%). This significant analysis of integrating all the possible coupling modes into a common framework and practical an integrated dynamic functional brain networks assists to explore the multiplexity of human brain dynamics under various conditions. These results untangled the swift of dominant coupling modes from α frequencies in awake condition to δ frequency in anesthesia (Purdon et al., 2013).

Analyzing neuromagnetic recordings from repeat scans under the same methodology revealed significant and reliable trends of complexity of brain activity and multiplexity of brain connectivity. Statistical analysis revealed a common frequency-dependent trend where DMN and FP demonstrate significant higher CI compared to the rest of brain networks across the frequency bands. Complementary, the CI was higher for the lower frequency bands (δ,θ,α1,α2) compared to the higher frequencies (β_1_,β_2_,γ). CI values were high reliable across the frequency bands and brain networks with ICC values above 0.85 (Fig.9.B).

FI values estimated within and between brain networks were also high reliable between scan session 1 and scan session 2. Interestingly, the more flexible set of pairs of brain networks are the DMN-CO, FP-CO following by DMN-CO and FP-S. It is worth to notice that the pattern of DICM is highly reproducible in both spatial scales whole network vs between networks. Our methodology harness the notion of DICM in order to reveal the multiplexity of human brain dynamics in both spatial scales using MEG resting-state activity. It is revealed that Δ and α_2_ frequencies govern the DICM and the multiplexity of neuromagnetic recordings at resting-state. The estimated ICC for both estimated comodulograms was 0.91 ±0.06 across the cohort.

A recent study revealed that higher functional brain dynamics between FP-DMN is correlated with higher cognitive flexibility (Douw et al., 2016).Here, we revealed that FP and DMN are the central core of brain flexibility in terms of dynamic reconfiguration of dominant coupling modes.

Finally, we demonstrated the proposed CI,FI and comodulograms in fMRI resting-state repeat scan recordings. The scan-averaged CI was estimated within thirteen brain networks and the related ICC. CI was statistically significant different between scale 1 and the rest 2-4 and also between scale 2 and scales 3-4 across brain networks. CI follows the frequency scales with the exception for scales 3 which had similar CI values with scale 4. ICC values for CI estimates were > 0.9 meaning that CI were high reliable.

This is the very first study that introduced the notion of cross-frequency coupling and DICM in fMRI. We found that the scan-averaged FI between and within the thirteen brain networks in both split-half scan sessions were high reliable with ICC values reaching the 0. 93 + 0.03 across scan – sessions.

Trial-averaged comodulogram across the whole brain network and between and within the thirteen brain networks reveal a significant trend where frequency scale 1 is the basic brain modulator following by scale 2. These pattern was observed also on the more detailed spatial scale of brain networks both within and between them. The methodology of DICM and FI will be useful to be applied to both cognitive tasks and disease cases.

It is important for any neuroscientist to understand the importance of these novel methodologies for understanding the complexity of brain activity (Antonakakis et al., 2016b; Dimitriadis et al., 2016b) and connectivity in functional neuroimaging. The incorporation of both intra and inter-frequency couplings to an integrated dynamic functional connectivity graph could reveal the pattern of DICM within and between brain networks and also their FI (Antonakakis et al., 2016a,2017a,b; Dimitriadis et al., 2015a,2015b,2016a,2016c,2017). Only with the adaptation of such methodology, the multiplexity of human brain connectivity could be revealed.

## 5. Conclusions

In the present study, we demonstrated a novel framework of exploring the complexity of brain activity and connectivity in the majority of functional neuroimaging modalities. The results were very promising for characterizing cognitive states at resting-state, during anesthesia, for designing reliable biomarkers and also for a better understanding of the multiplexity of functional brain connectivity. Present results further support previous findings focusing on dynamic reconfiguration of DICM as a framework of studying the brain rhythms and their possible interactions simultaneously. Multiplexity of human brain interactions can be explored only by integrating both intra and inter-frequency coupling modes into a brain network model.

## Acknowledgments

SID,DEL and KDS were supported by a MRC grant MR/K004360/1 (Behavioural and Neurophysiological Effects of Schizophrenia Risk Genes: A Multi-locus, Pathway Based Approach). This study received support from the UK MEG Partnership Grant (MRC/EPSRC, MR/K005464/1), CUBRIC and the School of Psychology at Cardiff University.

SID is also supported by a MARIE-CURIE COFUND EU-UK Research Fellowship. I would like to acknowledge Cardiff RCUK funding scheme for covering the publication fee.

## Author Contribution

Conception of the research analysis: SD; Methods and design: SD; Data analysis (SD); Drafting the manuscript: SD;

## Footnotes

1 http://neurotycho.org/spatial-map-ecog-array-task

## Abbreviations

DFBC –: dynamic functional brain connectivity
iPLV –: imaginary part of phase locking value
CFC –: cross frequency coupling
WC –: wavelet coherence
ROI –: regions of interest
EEG –: electroencephalography
MEG –: magnetoencephalography
FMRI –: functional magnetic resonance imaging
ROI –: Regions of Interests
DICM –: dominant intrinsic coupling modes
FI –: flexibility index
CI –: complexity index

